# Minimum Grid Cells for Place Cell Activity in Eleven Enclosures

**DOI:** 10.1101/2024.11.15.623794

**Authors:** Anu Aggarwal

## Abstract

The goal of this work was to uncover the minimum number of grid cells that can re-create the firing patterns of place cells in eleven different enclosures. This is useful for understanding firing patterns of place cells to design a very compact neurally-inspired spatial navigation system in hardware. For the project, we took place cell firing data from the Moser lab. Of the 342 place cells recorded, we selected one of the place cells firing in a single enclosure, one in six and one in all eleven enclosures. In MATLAB, we created grid cell firing patterns for 11250 grid cells. Connection weights between the place and grid cells were learned using machine learning-gradient descent algorithm. In all three cases, the place fields could be learned from at least 36 grid cells with three spatial firing frequencies, 2 directions for each frequency and 6 different phases for each direction. Including fewer phases reduced precision in deciphering the place. Weights learned were normally distributed with a wider spread and multimodal distribution for rooms with uneven, larger or multiple firing fields and narrower distribution otherwise. In rooms with no firing, all the weights varied a little around zero.

## 1. Introduction

Understanding the information processing in hippocampal formation has been one of the central questions in neuroscience ever since place-related firing patterns of hippocampal place cells were uncovered [1]. With place cell firing at the same place in the environment on repeated traversals of the animal through the same room, the hippocampal place cells are thought to encode for place in the environment in allocentric coordinates. To understand the origin of their unique firing patterns, other cells in the hippocampal formation were discovered. To mention a few, these are head direction cells (HDCs) and boundary vector cells (BVCs) in the subiculum [2][3][4], grid cells in the medial entorhinal cortex (MEC) [5], [6],[7], and cells in lateral entorhinal cortex (LEC). HDCs code for position of head, acting like compass for the brain. BVCs firing is proportional to the distance and direction from environmental boundaries. The grid cells fire repeatedly at corners of equilateral triangles in the environment creating a grid for the environmental map. The cells of the LEC are known to integrate object information [6]–[12].

Primarily, there are three categories of computational neuroscience models [13] explaining the place cell firing – the BVC based, the grid cell-based and the sensori-motor model. The BVC based model [14] considers place cell firing patterns to be due to the combination of inputs from BVCs. This is also known as the sensory model as BVC firing is thought to be determined solely by sensory inputs. This model was used to describe the observed place cell firing patterns very faithfully in [15]. The grid cell-based model [16], [7], [18], [19], [20] considers place cell firing to be the result of combination of inputs from several grid cells with different frequencies, phases and directions. The sensori-motor model [21],[22], and its proponents [23][24] consider place cell firing to be due to combination of inputs from both LEC and MEC. The MEC grid cells provide information about the place and LEC about the objects at the place which make it of significance. Thus, the place cell fires at a place that it finds significant enough to remember.

Anatomically [25]–[28], (**Fig. 1**) layers 2 of the LEC and MEC provide inputs to the DG, and CA3 and layers 3 to CA1 and subiculum via the perforant pathway. Layers 4 of the MEC and LEC are connected to each other. DG provides inputs to CA3 which provides inputs to CA1. Layers 5 and 6 of the LEC and MEC receive input from the CA1 and subiculum. CA1 provides inputs to subiculum which provides inputs to the post-rhinal and perirhinal areas. The post-rhinal and peri-rhinal areas also receive inputs from layers 5 and 6 of the LEC and MEC respectively and project back to layers 2 and 3 of the LEC and MEC. Thus, sensory inputs from the body to this region arrive at the entorhinal cortex (ERC) and subiculum (not the hippocampus). Outputs going to cingular cortex, amygdala, septal region, claustrum, infralimbic, association cortex and subcortical areas all go from layers 4-6 of the ERC (not hippocampus). The subiculum (which contains BVCs and HDCs) has bi-directional connections with the peri-, post-rhinal areas and the ERC. The ERC and subiculum both receive similar sensory inputs. We think that the subiculum uses this sensory information for HDCs and BVCs. And the grid cells use it for orienting to the cues. LEC and MEC are the only input to CA3 and DG and the CA1 receives inputs from LEC, MEC, and CA3 (not from subiculum). Spatial information is provided by the MEC and LEC provides object information, primarily. The grid cells integrate visual and other sensory information with motion, seen in grid cell remapping shown in [5], [18], [29], [30], [31], [32], [18] and present to the place cells which then fire specifically at a place defined by the grid cells providing inputs to the place cell. Another thing to note is that the grid cells project directly to CA3 and DG place cells without any synapses in between. Thus, connections between them can be modelled mono synaptically.

**Figure 1.**
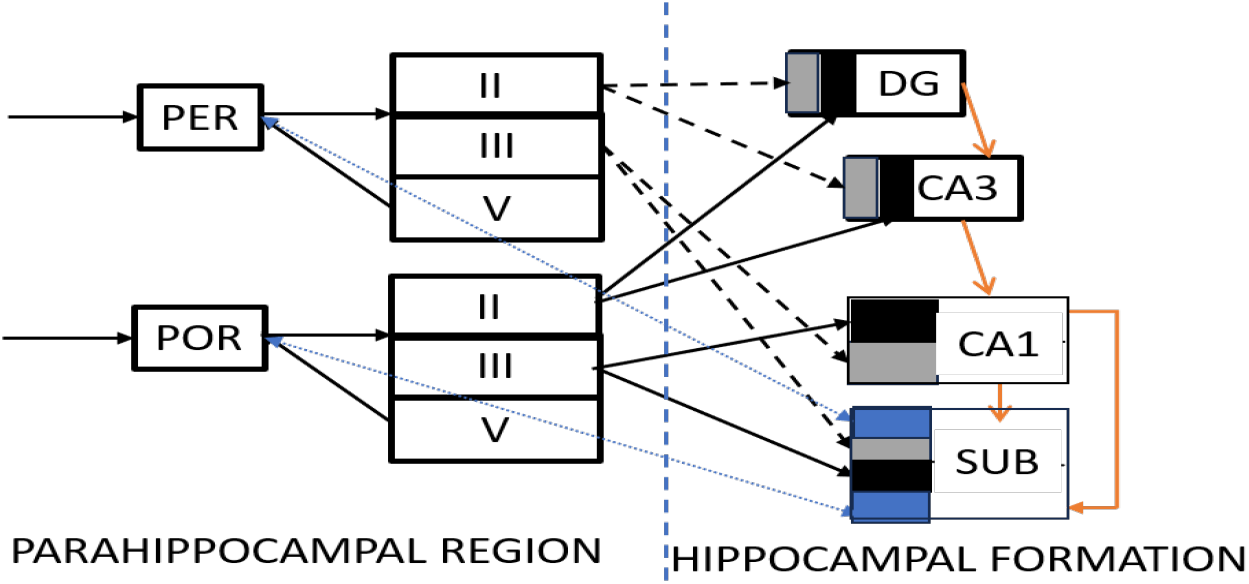
This figure is reproduced from [28]. It shows connections between rhinal cortex, entorhinal cortex, hippocampus and subiculum.

Anatomically, the BVCs have been found in the subiculum. But subiculum does not provide direct inputs to the place cells in the hippocampus. It projects back to MEC and LEC and hence, could impact the firing patterns of grid cells like change in direction with change in environment. Also, subiculum receives inputs from peri- and post-rhinal areas, therefore could be influenced by same sensory inputs as are received by the MEC and LEC. So, MEC grid cells could provide similar sensory inputs as BVCs to place cells. Moreover, it is harder to explain the place cell firing based on BVCs in bigger enclosures, especially in the center of the enclosure. The fidelity with which the BVC replicated the findings in [15] could be because firstly, in this study, only the place cell firings and not grid cell firings were recorded – therefore, it is harder to determine the impact of grid cells in place cell re-mapping. Also, grid cells might be receiving similar sensory inputs from post-rhinal areas as are received by BVCs which might cause similar changes in both with change in environmental geometry which might have led to misinterpretation of place cell firing being caused by BVCs instead of grid cells.

We think that the sensori-motor model also does not explain place cell firing as anatomically (Figure 1), there is separation of inputs from LEC and MEC throughout the hippocampal CA1, CA3 and DG. Also, the LEC cells represent information in egocentric coordinates while MEC and hippocampal place cells represent information in allocentric coordinates as also shown in [33].

Instead, the grid cell-based place cell firing model explains the place cell firing adequately but we think that it is not a motor or path integration model alone [34]. We surmise that the grid cells integrate both the motor and sensory inputs and present them to the place cells as also mentioned in [35]. Indications of this come to some extent from studies like [30], [36]. The grid cells are arranged in modules [16], [5], [19], [36], [20], [37]. Grid cells in a module share a frequency, 2 directions separated by 30 degrees and have several phases that map the environment. The grid cell frequencies scale by 1.41 along the dorso-ventral axis. There are around 4-10 modules per animal. Grid cells have been shown to undergo re-direction or translation with change in environments and anchoring cues within the environment even when the color, size and shape are the same but location of entrance and of orienting cues is different [5], [18], [29], [30], [31], [32], [36], [38]. This is thought to be the reason for place cell re-mapping seen in a new environment. Since there are multiple grid cells in a module with different phases and each grid cell has several fields, several grid cells from a module can be firing at the same location. In this work, we wanted to find out the minimum number of grid cells that could replicate place cell firing patterns. This is useful to solve a neuroscience problem and for designing compact silicon VLSI chip-based models of the hippocampus for use in spatial navigation systems. The smaller and more compact the system, more economical it will be. Therefore, in this project we took 342 place cell recordings from CA3 of 8 animals in eleven different rooms from the Moser lab [39]. We created 11250 grid cell firing patterns in Matlab using the model described in [40]. As seen anatomically (Figure 1), the projection from grid cells in layer 2 and place cells in CA3 is direct and hence, the distance between the two is single synapse and gradient descent algorithm can be used to learn the place cell firing patterns as seen in Moser lab data from grid cells that we created in Matlab. Of the 342 place cells, we took 3 representative place cells. From studies like [30],[38], plasticity seems to be guiding the learning between grid and place cells in different enclosures. Hence, we used the gradient descent learning algorithm which is mathematical but like natural learning in brain.

## 2. Methods

The lab data [39] contained recordings from 342 CA3 place cells from 8 animals in 11 different rooms. With an activity threshold of 0.1Hz, around 210 cells were active in at least one room, 103 were active in only one room, 44 were active in two rooms, 21 were active in six or more rooms, 10 were active in eight or more rooms and 2 of the cells in the distal most CA3 fired in all 11 rooms. As mentioned earlier, we took three of these place cells for further analysis-the first one fired in all the 11 rooms, the second one in 6 rooms and the third one in only 1 room. We created grid cell firing patterns with three different spatial firing frequencies (28, 39, 55), 6 different directions (0, 10, 20, 30, 40, 50 degrees) and 625 different phases (25 in the x and 25 in the y-direction) in a 1m x 1m enclosure based on the model described in [40]. The firing frequencies were 1.41 times the previous one as reported in [16]. We used only 3 frequencies (28, 39, 55 cm) for our project as the place cell recordings were from the dorsal part of the CA3 corresponding to which we expect grid modules of only the lowest frequencies. We used grid cells with only 2 directions (out of the 6 created in Matlab) that were 30 degrees apart for each frequency as reported in [16]. Grid cell directions were varied with the frequency and the room that the animal was in as seen in [5], [18], [29], [30], [31], [32]. We took six phases for each frequency and direction to run our gradient descent learning algorithm and learn the observed place cell firing pattern.

Initially, we connected the first cell (firing in all the 11 rooms), to all 11250 grid cells with random weights. Then, we trained the weights using gradient descent algorithm to learn the firing patterns of place cells as seen in the lab. We used 100 trials of 10,000 iterations each of the gradient descent algorithm with a learning rate of 0.1. We analyzed the learned weights for maximum weights for each of the 3 frequencies in each of the 11 rooms. We took the maximum weight direction and the direction roughly 30 degrees away from it for each room. Then, we also took 5 phases surrounding the phase corresponding to maximum weight. And we re-ran the gradient descent to learn the place cell firing patterns again. Since there are overlaps in the modules [16] and each module has around two different directions 30 degrees apart and all possible phases, this selection of grid cells is biologically plausible. To determine the minimum number of grid cells that could replicate the place cell firing pattern, we kept reducing the number of phases from 6 to 5 to 4 to 3 to 2 to 1 and re-ran the gradient descent for each of the cases. Thereafter, we reduced the number of directions from 2 to 1 while keeping frequencies at 3 and phases at 6 and re-ran the gradient descent algorithm. Next, we re-ran the gradient descent algorithm with grid cells of different number of frequencies (1 for each module), 2 directions per module and 6 phases. We tried different combinations of grid cells to determine the minimum number of grid cells that could successfully replicate the place cell firing patterns. Same method was repeated for selection of grid cells connected to place cells for the second and third place cells - firing in 6 out of 11 and 1 out of 11 enclosures. Subsequently, all sets of weights obtained for the combinations of grid cells were analyzed to understand the connectivity. For this, the histogram of weights was created for each of the eleven rooms for 100 trials of each experiment. The magnitude of weights was also analyzed for each direction, phase and frequency of the grid cell that is connected to the place cell.

## 3. Results

### A. Experiment 1

In the first experiment, one of the place cells firing in all eleven rooms (Fig. 1(a)) was connected to all 11250 grid cells with three (28, 39 and 55cm) spatial firing frequencies, 625 phases and 6 directions. The results are shown in Fig. 2 (b) and compared with the recorded place cell firings in Fig 2(a). The firing patterns in all the 11 rooms could be learned successfully. In Figs 2(a) and (b), the room numbers go from left to right in each row (1→ 4) and then in the next row (5→ 8 and 9→11) in increasing order. Next, we analyzed the weights from the 100 trials. Firstly, we plotted histograms of mean weights for each room (Fig. 2 (c)). Weights learned were normally distributed. The spread of weights is wider for rooms with uneven, larger or multiple firing fields. While for the rooms with smaller firing fields, the weight distribution is narrower, and for those with medium sized fields, weight distribution has a medium spread. For small and medium firing fields, the weights have unimodal distribution. Larger and multiple firing fields are associated with multimodal distribution of weights. Weights did not differ a lot across the 100 learning trials. But weights learned for each room exhibited a different pattern. To determine how much each grid cell with a given phase, direction and spatial frequency contributes to a place cell firing, we plotted mean weights versus phase and direction. The results from first direction and 625 phases are shown in figure 2(d) as representative of all 11250 grid cells. Here, the phases are marked along the x-axis, for first direction. The weights vary sinusoidally with the phase and direction for a frequency. Rooms with small single place firing patterns (rooms 1, 3, 9 and 11) are associated with similar and smaller contributions from each grid cell with different spatial frequency. Rooms with relatively bigger but regular single place fields (rooms 2, 4, 8 and 10) have higher contribution from grid cells with higher spatial firing frequencies (39 and 55cm) rather than grid cells with smaller spatial firing frequency (28cm). In room number 7, the place field is big and irregular. If we examine the weight distribution in this room, it has higher contribution from the 55cm spatial frequency. The weights associated with this frequency have multiple waves of overlapping sinusoids.

**Figure 2.**
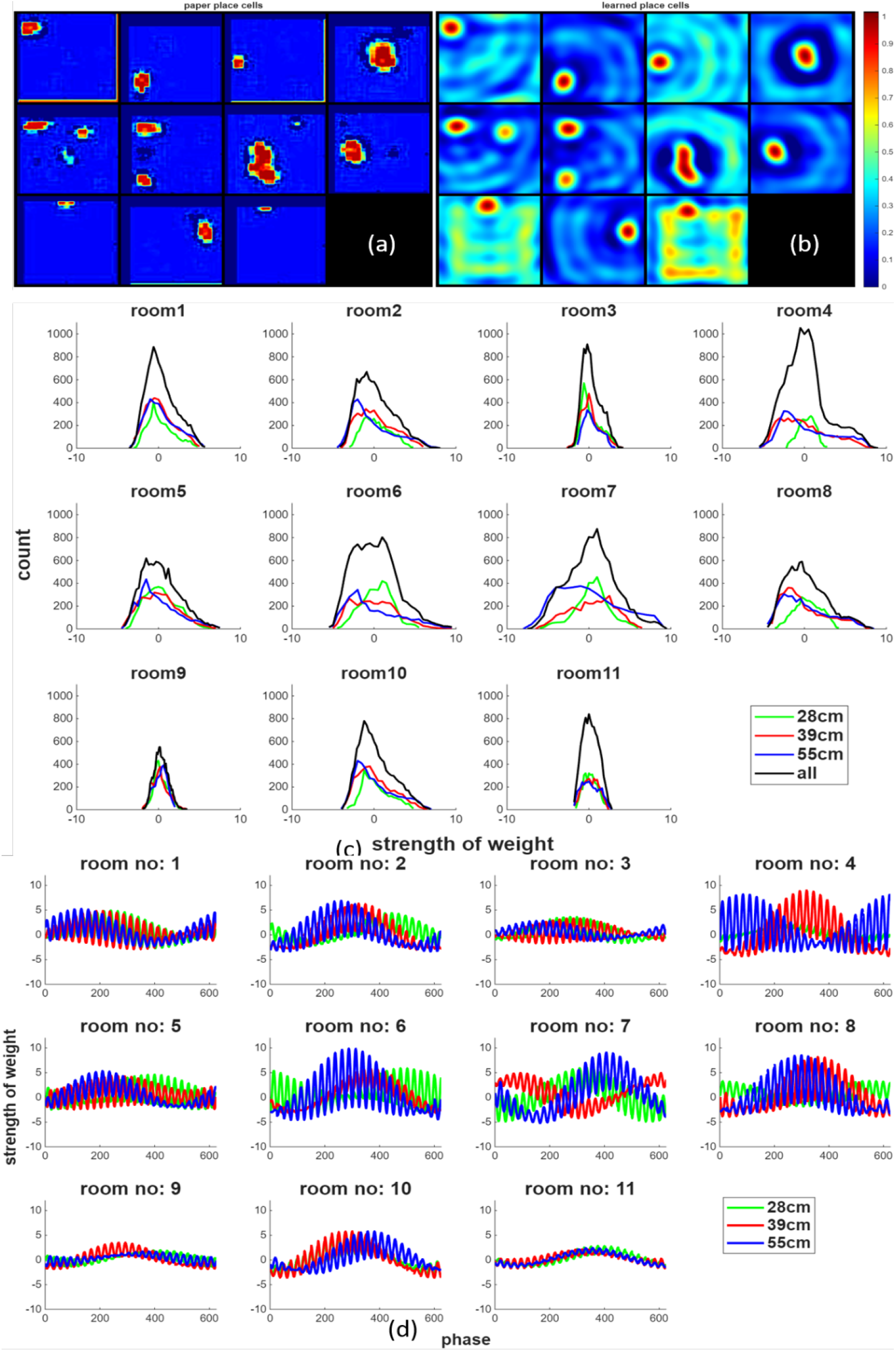
For (a) and (b), room numbers are labelled from 1 to 4 horizontally in the first row in that order, 5-8 in the second row and 9-11 in the third row. (a) recorded place cell firings in eleven different rooms. (b) learned place cell firing patterns in the eleven rooms using gradient descent learning algorithm, 11250 grid cells with 625 phases, 6 directions and 3 frequencies. It is seen that all place cell firing patterns are learnt successfully. (c) all learned weights over the 100 trials with 10,000 iterations each have a gaussian distribution. The weights for rooms with smaller firing patterns have a narrower distribution than the ones with a larger firing pattern. The peak of the distribution is centered around 0. (d) phase label on the x-axis indicates phase of firing of the grid cell. Results are shown for a single direction and 3 frequencies for clarity. The strength of the weights varies roughly sinusoidally with the phase and direction of the grid cell. The pattern varies with the room.

### B. Experiment 2

To determine the minimum number of grid cells that can replicate the place cell patterns observed in all the enclosures, we kept on reducing the number of grid cell inputs while keeping the grid cell combination biologically plausible as seen in [16]. We started with grid cells of all three frequencies, two directions and six phases. Subsequently, we reduced the number of phases from 6→ 4 → 3 → 2 →1. We could see the place cell firing patterns (Fig. 2b) being replicated with grid cells of 6 phases at least, i.e., 36 grid cells in total (3 frequencies, 2 directions for each frequency and 6 phases for each direction (Table 1)). Then we reduced the number of frequencies from 3 → 2 →1 while keeping the number of directions at 2 and phases at 6. In the next trials, we reduced the number of directions while keeping the number of frequencies at 3 and phases at 6. None of the other combinations could replicate the grid cell firing patterns completely as shown in Fig. 3b.

**Table 1.**
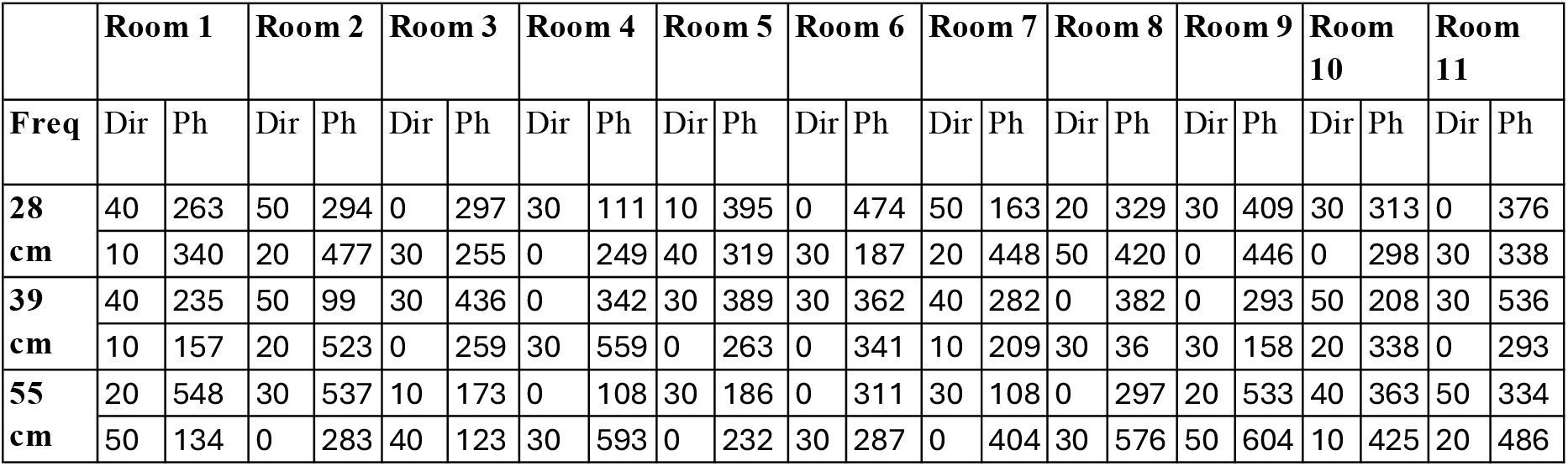
showing frequency, direction and starting phase of grid cells used in the second experiment. Next 6 phases from the starting phase were included in the simulations.

|  | Room 1 |  | Room 2 |  | Room 3 |  | Room 4 |  | Room 5 |  | Room 6 |  | Room 7 |  | Room 8 |  | Room 9 |  | Room 10 |  | Room 11 |  |
| --- | --- | --- | --- | --- | --- | --- | --- | --- | --- | --- | --- | --- | --- | --- | --- | --- | --- | --- | --- | --- | --- | --- |
| Freq | Dir | Ph | Dir | Ph | Dir | Ph | Dir | Ph | Dir | Ph | Dir | Ph | Dir | Ph | Dir | Ph | Dir | Ph | Dir | Ph | Dir | Ph |
| 28<br>cm | 40 | 263 | 50 | 294 | 0 | 297 | 30 | 111 | 10 | 395 | 0 | 474 | 50 | 163 | 20 | 329 | 30 | 409 | 30 | 313 | 0 | 376 |
|  | 10 | 340 | 20 | 477 | 30 | 255 | 0 | 249 | 40 | 319 | 30 | 187 | 20 | 448 | 50 | 420 | 0 | 446 | 0 | 298 | 30 | 338 |
| 39<br>cm | 40 | 235 | 50 | 99 | 30 | 436 | 0 | 342 | 30 | 389 | 30 | 362 | 40 | 282 | 0 | 382 | 0 | 293 | 50 | 208 | 30 | 536 |
|  | 10 | 157 | 20 | 523 | 0 | 259 | 30 | 559 | 0 | 263 | 0 | 341 | 10 | 209 | 30 | 36 | 30 | 158 | 20 | 338 | 0 | 293 |
| 55<br>cm | 20 | 548 | 30 | 537 | 10 | 173 | 0 | 108 | 30 | 186 | 0 | 311 | 30 | 108 | 0 | 297 | 20 | 533 | 40 | 363 | 50 | 334 |
|  | 50 | 134 | 0 | 283 | 40 | 123 | 30 | 593 | 0 | 232 | 30 | 287 | 0 | 404 | 30 | 576 | 50 | 604 | 10 | 425 | 20 | 486 |

**Figure 3.**
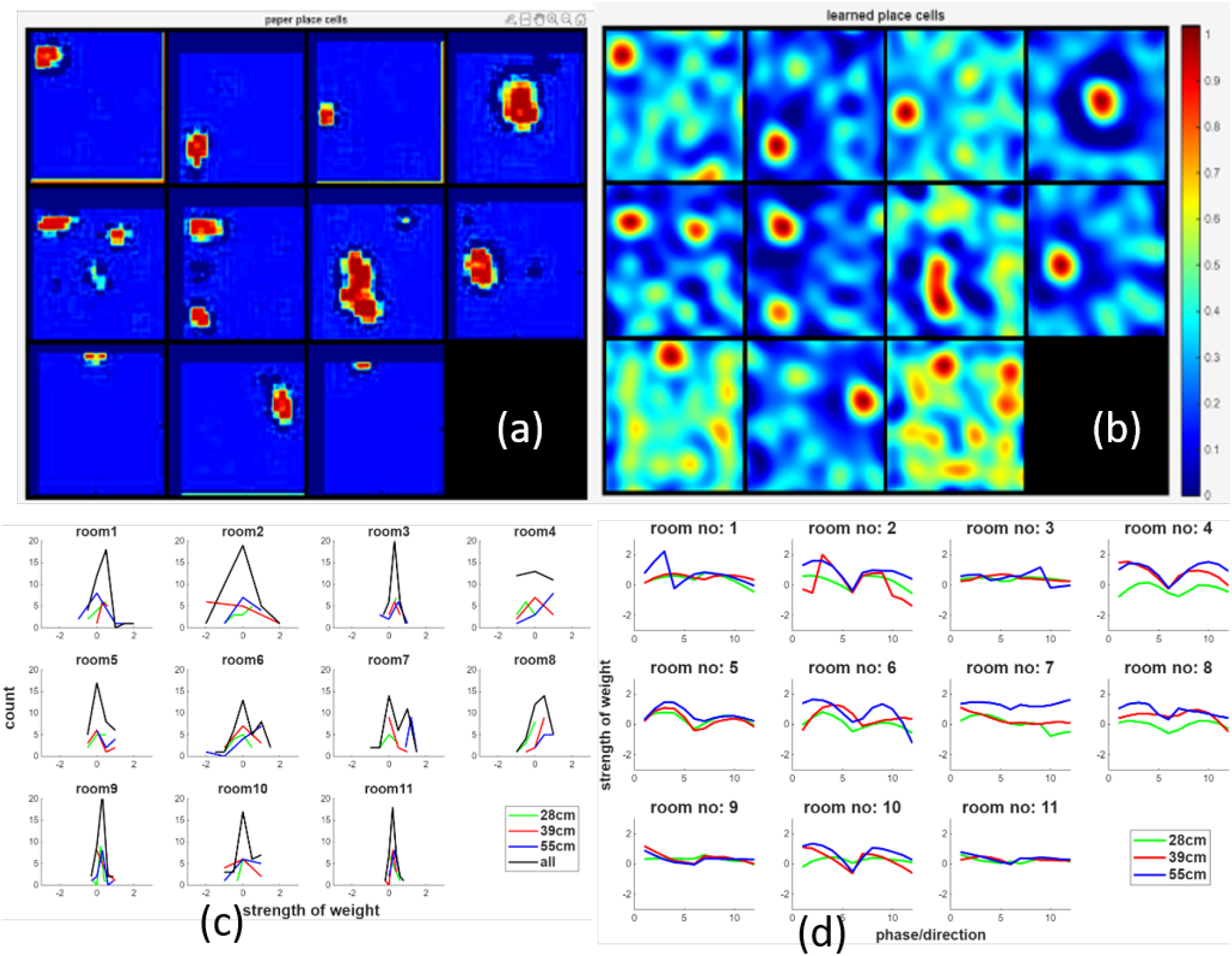
For (a) and (b), room numbers are labelled from 1 to 4 horizontally in the first row in that order, 5-8 in the second row and 9-11 in the third row. (a) recorded place cell firings in 11 different rooms. (b) learned place cell firing patterns from 36grid cells with three different spatial firing frequencies (28, 39 and 55cm), 2 directions and 6 phases. The results indicate that the unique firing patterns of the place cell could be learnt for each room separately with inputs from only 36grid cells. (c) Histograms of mean weights over 100 trials indicate that the weights have normal distribution. Rooms with smaller place fields (rooms 1, 3, 9 and 11) have narrower and those with bigger fields have wider distributions. Rooms with multiple or irregular place fields (rooms 5, 6 and 7) have multimodal distribution of weights while those with single place fields have unimodal distribution. (d) shows sinusoidal variation in magnitude of weights with phase and direction. X-axis indicates 6 phases for each of the 2 directions for the 3 frequencies. Rooms with bigger fields have higher contribution from the grid cells with higher spatial frequencies. Rooms with small single place firing patterns (rooms 1, 3, 9 and 11) are associated with similar and smaller contributions from each grid cell with different spatial frequency. Rooms with relatively bigger but regular single place fields (rooms 2, 4, 8 and 10) have higher contribution from grid cells with higher spatial firing frequencies (39 and 50cm) rather than grid cells with smaller spatial firing frequency (28cm). In room number 7, the place field is big and irregular and has a higher contribution from the 55cm spatial frequency.

The weights for the case with 36 grid cells (3 frequencies, 2 directions and 6 phases) where place cells were learned were analyzed next. Histograms of weight distribution from mean of 100 trials can be seen in Fig. 3 (c). It shows a normal distribution of the weights for all rooms. For each room, the weight distribution is different than that for other rooms. Analyzing the magnitude of weights for different directions and phases in each room, there is sinusoidal pattern (Fig. 3(d)). It is seen in all the rooms though every room has a different phase and direction of grid cells. Here, the x-axis shows the grid cell number with 2 directions and 6 chosen phases (Table 1) in that order. First 6 indices denote the 6 phases of first direction, and next 6 indices denote the 6 phases of the second direction for each frequency. Grid cells with different frequencies are shown in different colors.

#### C. Experiment 3

For this experiment, we studied a place cell which fired in 6 of the eleven rooms (Figure 4(a)). Following the procedure of Experiment 2, we could replicate the place cell firing patterns (Figure 4b) when at least 36 grid cells of 3 frequencies, 2 directions and 6 phases were connected to the place cell. Next, we created histograms of the mean weights over the 100 trials (Figure 4c). Similar normal distribution of weights was observed as for the cell firing in all 11 rooms except for a narrower distribution. The mean was centered around 0. The spread of the distribution was more for the rooms with firing (rooms 1, 2, 3, 6, 8, 9) than without firing. The spread was also more for rooms with bigger and irregular (room 1, 2, 6, 8, 9) place fields than those with smaller fields (room3). No multimodal distribution was observed for any room. Even though distributions are similar for rooms with no firing, they are different from each other in rooms where the place cell fires (rooms 1, 2, 3, 6, 8, 9). Next, distribution of weight magnitudes for different phase and direction of grid cells was plotted (Figure 4d). The distribution has sinusoidal variation as seen in Figure 2(d) but the magnitude is lower. For the rooms with no place cell firing (rooms 4, 5, 7, 10 and 11), the contribution from all the grid cells was similar irrespective of frequency. For rooms with increasing size of place fields (rooms 3, 2, 8, 9 and 1 in that order), the magnitude of weights for various grid cell frequencies increases. For rooms with irregular place cell firing pattern (room 6), the maximum contribution is from grid cells with higher spatial firing frequencies.

**Figure 4.**
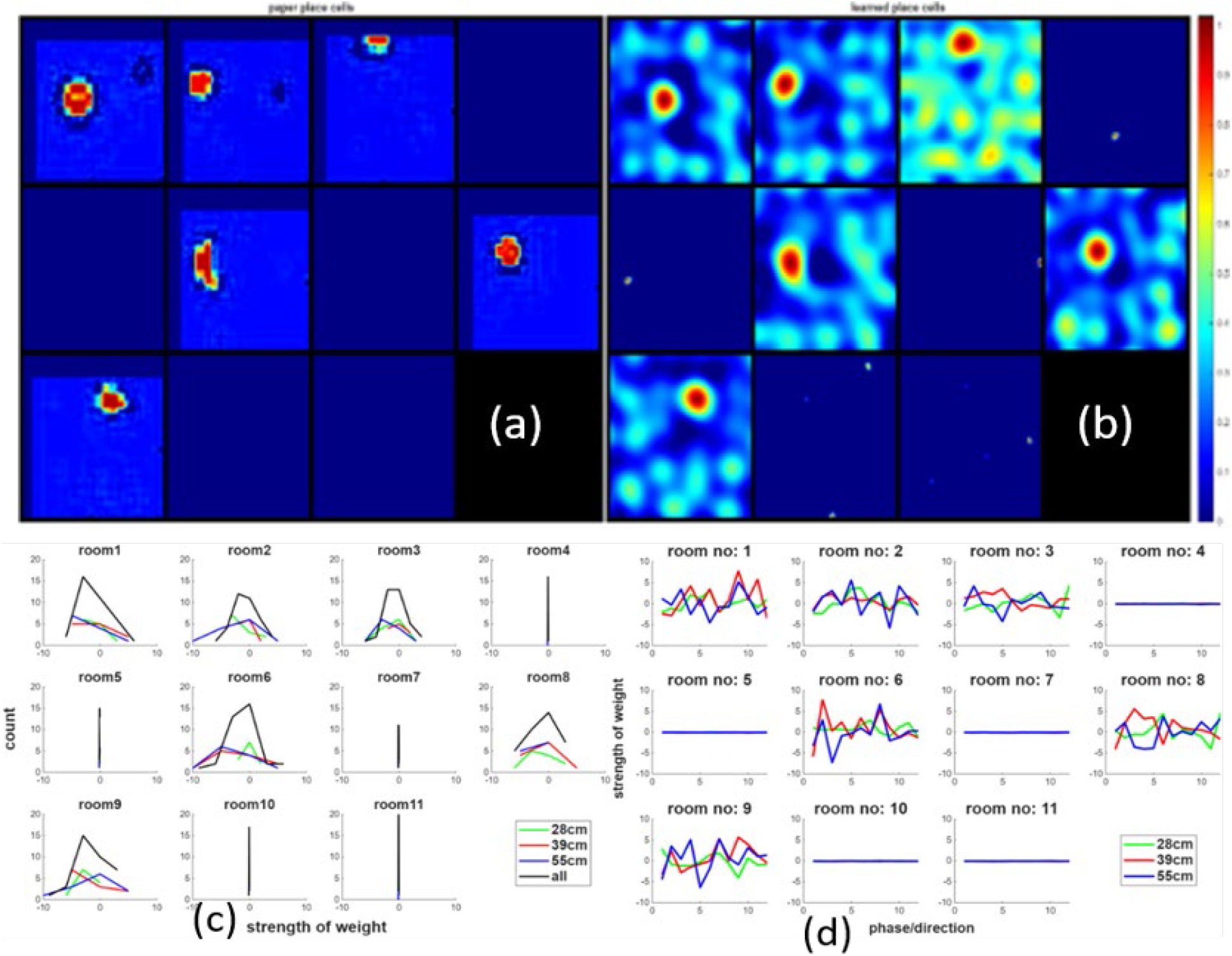
For (a) and (b), room numbers are labelled from 1 to 4 horizontally in the first row in that order, 5-8 in the second row and 9-11 in the third row. (a) Recorded place cells in eleven different enclosures. (b) Successfully learned place cell firing patterns in the eleven rooms with 36 grid cell inputs using gradient descent algorithm. (c) Histogram of mean weights learned over 100 trials shows the normal distribution of weights. The weights corresponding to rooms with bigger firing patterns have a wider distribution than those corresponding to smaller or no place fields (rooms 3, 4, 5, 7, 10, 11). No multimodal distribution is seen for any room. Even though distributions are similar for rooms with no firing, they are different from rooms where the place cell fires (rooms 1, 2, 3, 6, 8, 9). (d) x-axis labels indicate 6 grid cell phases for 2 directions. For rooms with smaller or no place fields (rooms 3, 4, 5, 7, 10, 11), there is small equal contribution from grid cells with all frequencies. For rooms with bigger firing patterns, the contribution from grid cells with all spatial frequencies increases with maximum contribution from higher frequencies.

### D. Experiment 4

Lastly, we wanted to explore the grid cells to place cell connectivity for the 103 place cells that fired in a single room. We learned the connection weights of all these cells (taken one at a time) using the techniques described above. Here, we present results from one of the representative cells. The recorded and learned place cells are shown in Figure 5(a) and (b) respectively. These place cell firing patterns could be learned by using at least 36 grid cells of 3 frequencies, 2 directions and 6 phases. Corresponding weights learned are shown in Figure 5(c) and (d). From Figure 5(c), we can see that all the weights are normally distributed. The weights in the room where the place cell is firing are more widespread while those in the rooms where the place cell is not firing are more narrowly distributed. This is like what has been observed in previous experiments. In figure 5(d), weight distribution with phase and direction for a frequency shows sinusoidal distribution of the weights for all rooms with similar magnitude of weights for all rooms except for room 5 where the place cell is firing.

**Figure 5.**
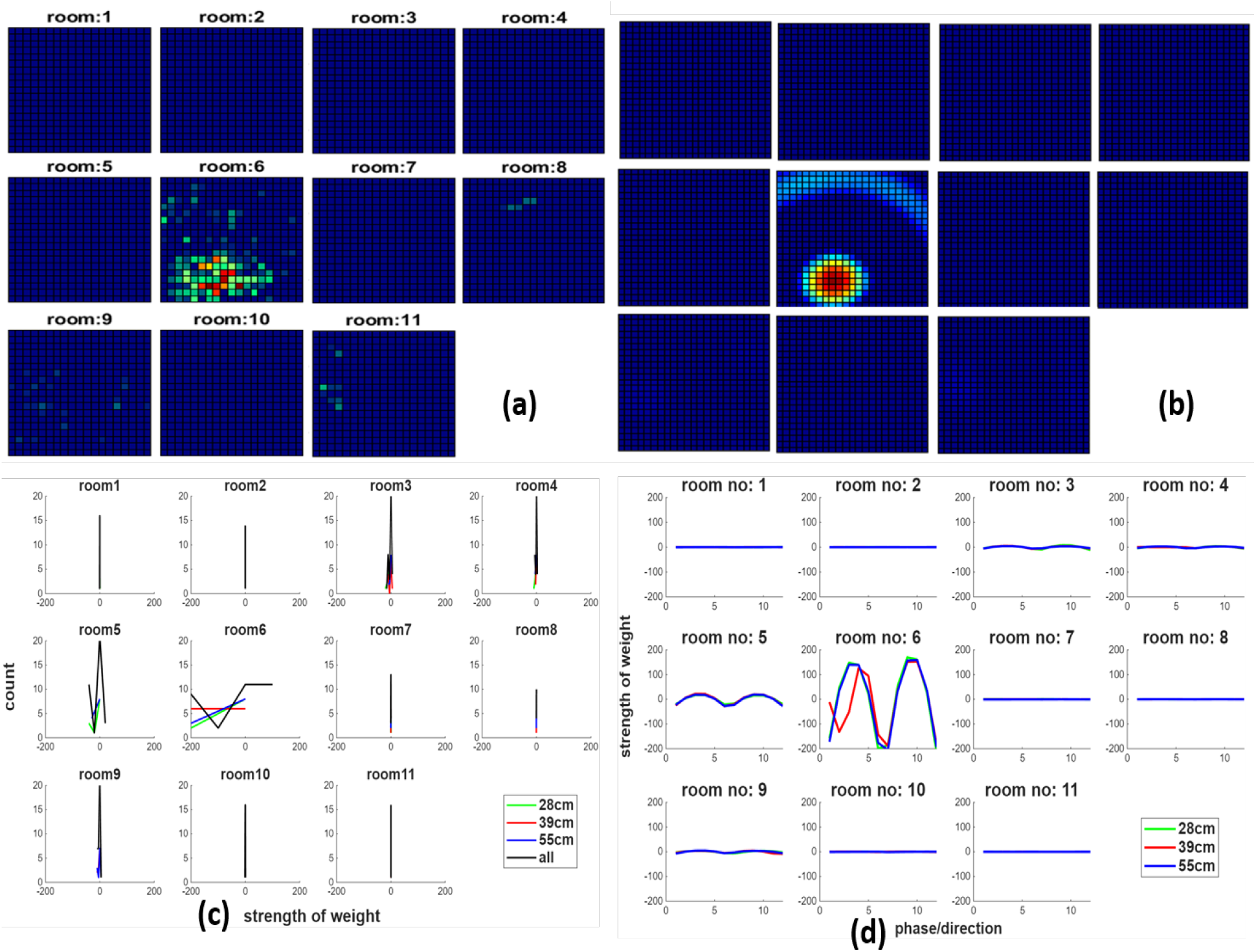
For (a) and (b), room numbers are labelled from 1 to 4 horizontally in the first row in that order, 5-8 in the second row and 9-11 in the third row. (a) place cell recording from eleven rooms shows the place cell firing in room 5. This was successfully replicated by learning weights using the gradient descent learning algorithm, as shown in (b). (c) Histograms of weights from 100 trials are similar to those seen in Figs. 1-3 but much lower in magnitude. Most of the rooms have narrow weight distribution, peaking around zero. Only in room 5, there is a wider distribution of weights which is off-centered from 0. Weights from different frequencies have different peaks. (d) x-axis labels indicate grid cell phases from 1 to 12 with first 6 belonging to the first direction and the next 6 to the next direction. On plotting the magnitude of weights against direction and phase of grid cells, for all three freqeuncies of grid cells separately, it is evident that the pattern is sinusoidal as seen in prior experiments, contributions from all frequencies is similar in room 5 where the place cell is firing.

## 4. Discussion

The goal of this work was to uncover the minimum number of grid cells that can replicate the firing patterns of 342 place cells recorded in eleven different enclosures. As a representative of the population, place cells which were firing in either all eleven enclosures, six enclosures or a single enclosure were included in the analysis. In all the experiments, the place cell firing patterns were successfully learnt using 36 grid cells. From our results, we think that at least 36 grid cells with at least 6 different directions (2 for each frequency) are required to re-create any of the observed firing patterns. If we choose fewer grid cells as have been observed experimentally, i.e., modular arrangement, then the resulting place cell firing pattern is less precise. Alternatively, if we take grid cells of a single frequency but several directions, then fewer grid cells are required to recreate the firing patterns. However, the latter arrangement is not seen biologically in lab experiments, hence, we do not report our results from such simulations here.

Our results indicate that grid cells with two directions separated by 30 degrees in a module are involved in place cell firing. Also, multiple phases are involved in firing – meaning that the place cells provide code for a position in space as defined by several grid cells and not just one. For the same place cell, there is a different firing pattern in different rooms. This firing pattern is achieved when the grid cell that it is connected to changes its direction and phase corresponding to the orienting cues. This is remapping at the level of grid cells. This also seems to be justified by [42] where there was change in direction of grid cells as the animal moved to another environment. This reorientation of grid cells in different rooms can explain how the same connectivity between the grid cells and the place cell can cause the re-mapping of place cell due to re-mapping of grid cells in different rooms.

Further, there seems to be correspondence between frequency of grid cells and firing field size of the connected place cell. In the rooms with smaller fields, contribution of grid cells with smaller frequencies is more than in rooms with larger or more irregular fields. On analyzing the connection weights between the grid and place cells, there was a unimodal distribution of weights for single place fields and multimodal distribution for multiple, bigger or irregular firing fields. This indicates that the small, single place fields can be realized with single grid cell spatial frequency whereas multiple, bigger or irregular place fields require multiple grid cell frequencies. The sinusoidal distribution of weights along the phase and direction of grid cells is expected given the periodic firing patterns of grid cells.

We observe that for a place cell firing in any number of enclosures, we need at least 36 grid cells to fire at the same time. This is very similar to observations in [43]. Even though the observations in [43]were for single place fields for each place cell as opposed to current study where we took 3 place cells with different number of place fields in different enclosures. The 1000 place cells used in [43] were generated computationally, whereas in our study, the 342 place cells were taken from lab recordings from 8 animals in 11 different enclosures. That study also considered single place fields in an enclosure 4mX4m whereas our study took lab data from firings in 1mX1m enclosures due to which there were multiple and/or irregular place fields in some of the enclosures.

We observed that more than frequency, direction plays a role in changing the number of grid cells that could generate the place cell firing patterns successfully in all 3 cases. We need at least 6 different directions – 2 for each frequency to achieve it as biologically, a frequency of grid cells corresponds to a direction and all grid cells in a module share a frequency and 2 directions. If there were grid cell modules with different directions but same frequency, then we could achieve place cell firing patterns with fewer grid cells than observed. For instance, in one of the experiments, we saw single room place cell firing patterns could be replicated with 18 grid cells of single frequency, 6 directions and 3 phases. Based on the anatomy [25], [26] and our results, we think that the place cells could be realized at the corresponding places in the hippocampus where several modules of grid cells co-exist [42]. So that grid cells of different frequencies could be connected to single place cells.

From all the above, it is surmised that the grid cells resolve the space into spatial distance, direction and phase and combine sensory and motor inputs and present to place cells. Each grid cell fires at multiple locations. Thus, several grid cells in a module with different phases could be firing at the same time which corresponds to animal position at a location. So, each place cell firing can be a result of combination of firings of several grid cells at the same time. Also, several different places can be coded by the same grid cell when combined with firings from other grid cells. In other words, animal motion and sensory inputs generate grid cell firings which in turn code for several different places in the environment through place cell firing in the environment. For instance, if the grid cells of three frequencies, 2 directions (30 degrees apart) and 6 adjoining phases are included in the firing of a place cell in say, room1, it means that the code for the place is defined by a combination of 3 frequencies – 28, 39 and 55 cms, 2 directions (30 degrees apart) and 6 phases (mentioned in Table 1).

We surmise that since place cell firing happens at a phase of the theta [44], ensembles of place cells, firing at different phases of theta can possibly store memories for an episode in pre-frontal cortex in a sequence determined by the theta phase (Figure 6). This not only stores the position defined by the place cell from grid cell inputs, but it will also encode LEC processed object information. But retrieval process uses the place cell as an index to retrieve the entire context stored at the memory location. If memories are stored in a Hopfield network, then each place cell acts like an index to a network with adjoining networks tied to each other with theta phase. This builds episodic memories as a network of several Hopfield networks. The information about place and object related information from the LEC is stored in memories in the pre-frontal cortex with index provided by the place cell. Whenever the place cell is re-activated, it helps retrieve all the LEC and MEC information encoded in the memories in the pre-frontal cortex in the correct sequence. Thus, place cell acts like an index to the event and all the place cells encoding an event are tied in a temporal chain to replay the event as required.

**Figure 6.**
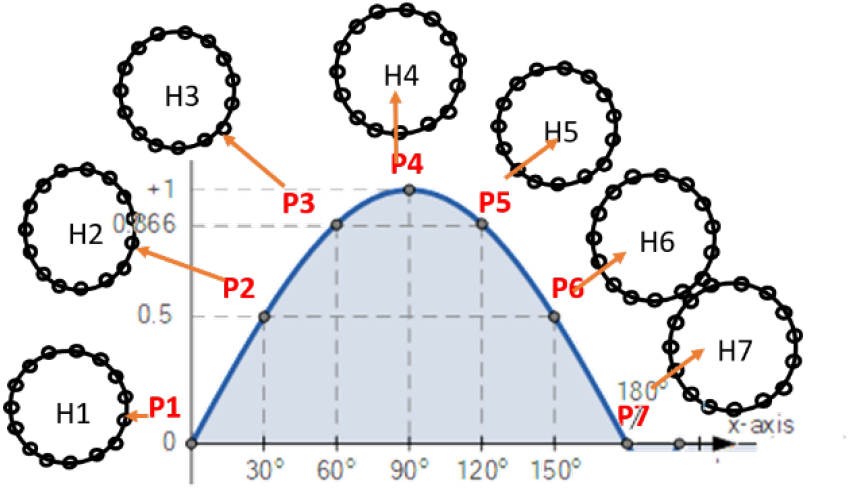
Cartoon representation of possible weaving of memories with theta phase. X-axis is the phase of theta and circles represent the Hopfield networks.

## 5. Conclusion & future work

We conclude that each place cell is connected to several grid cells from at least 3 modules which means grid cells with 3 different frequencies and 2 different directions. And each place cell firing needs connection to at least 36 grid cells. If fewer grid cells (fewer phases, directions or frequencies) are connected to place cell, its precision decreases. The grid cell remapping underlies the place cell remapping seen in cases where the place cell fires in multiple environments. Future work involves developing this system in silicon based on the results presented here and use it for spatial navigation, path planning and episodic memory storage.

## 6. Acknowledgements

The author thanks the Moser lab for very kindly sharing the data that was analyzed in this paper. A big thanks to Prof Edward I Moser and Prof Michael E. Hasselmo for providing invaluable feedback on the work during the SFN, 2024 and later.

## References

[1] J. O’Keefe, & J. Dostrovsky, 1971, “The hippocampus as a spatial map-Preliminary evidence from unit activity in the freely moving rat”, Brain Res, 34 (1), 171–175.

[2] J. S. Taube, 1998, “Head direction cells and the neurophysiological basis for a sense of direction”, Prog Neurobiol. 1998 Jun; 55(3):225–56. doi: 10.1016/s0301-0082(98)00004-5. PMID: 9643555.

[3] E. J. Golob, J. S. Taube, 1997, “Head direction cells and episodic spatial information in rats without a hippocampus”, Proc Natl Acad Sci U S A. 1997;94(14):7645–7650. doi:10.1073/pnas.94.14.7645.

[4] C. Lever, S. Burton, A. Jeewajee, J. O’Keefe, N. Burgess, 2009, “Boundary vector cells in the subiculum of the hippocampal formation”, J Neurosci. 2009 Aug 5;29(31):9771–7. doi: 10.1523/JNEUROSCI.1319-09.2009. PMID: 19657030; PMCID: PMC2736390.

[5] T. Hafting, M. Fyhn, S. Molden, M. B. Moser., E. I. Moser, August 2005, “Microstructure of a spatial map in the entorhinal cortex”, Nature, 436, 801–806.

[6] M. Hasselmo, M. P. Brandon, 2012, “A model combining oscillations and attractor dynamics for generation of grid cell firing”, Frontiers in Neural Circuits, Vol 6, Article 30, pp 1–13.

[7] B. L. McNaughton, F. P. Battaglia, O. Jensen, E. I. Moser & M. B. Moser, 2006, “Path integration and the neural basis of the ‘cognitive map’”, Nature Reviews Neuroscience, 7, 663–678.

[8] B. J. Young, T. Otto, G. D. Fox, H. Eichenbaum, 1997, “Memory representation within the parahippocampal region”, J. Neurosci. 17, 5183–5195.

[9] E. L. Hargreaves, G. Rao, I. Lee, J.J. Knierim, 2005, “Major dissociation between medial and lateral entorhinal input to dorsal hippocampus”, Science 308, 1792–1794.

[10] D. Yoganarasimha, G. Rao, J.J. Knierim, 2011, “Lateral entorhinal neurons are not spatially selective in cue-rich environments”, Hippocampus 21, 1363–1374.

[11] M. Pilkiw, et al., 2017, “Phasic and tonic neuron ensemble codes for stimulus-environment conjunctions in the lateral entorhinal cortex”, eLife Sci. 6, e28611.

[12] S. S. Deshmukh, J.J. Knierim, 2011, “Representation of non-spatial and spatial information in the lateral entorhinal cortex”, Front Behav. Neurosci. 5, 69 (2011).

[13] E. Moser, Y. Roudi, M. Witter, et al., 2014, “Grid cells and cortical representation”, Nat Rev Neurosci 15, 466–481 (2014). 10.1038/nrn3766.

[14] R. M. Grieves, E. Duvelle, P. A. Dudchenko, 2018, “A boundary vector cell model of place field repetition”, Spatial Cognition and Computation, 18(3), 217–256.

[15] Roddy M. Grieves et al., 2026, “Hippocampal place cells map terrain geometry independently of behavior”, Sci. Adv. 12, eadz9893(2026). DOI: 10.1126/sciadv.adz9893

[16] H. Stensola, et. al., 2012, “The entorhinal grid map is discretized”, Nature, 492, 72–78.

[17] N. Burgess, C. Barry, J. O’Keefe, 2007, “An oscillatory interference model of grid cell firing”, Hippocampus, 17, 801–812.

[18] M Fyhn, T Hafting, A Treves, MB Moser, EI Moser, 2007, “Hippocampal remapping and grid realignment in entorhinal cortex”, Nature, 8;446(7132):190–4. doi: 10.1038/nature05601. Epub 2007 Feb 25. PMID: 17322902.

[19] Y. Gu, S. Lewallen, A. A. Kinkhabwala, C. Domnisoru, K. Yoon, J. L. Gauthier, I. R. Fiete, D. W. Tank, 2018, “A Map-like Micro-Organization of Grid Cells in the Medial Entorhinal Cortex”, Cell, 175(3):736–750.e30. doi: 10.1016/j.cell.2018.08.066. Epub 2018 Sep 27. PMID: 30270041; PMCID: PMC6591153.

[20] J. G. Heys, K. V. Rangarajan, D. A. Dombeck, 2014, “The functional micro-organization of grid cells revealed by cellular-resolution imaging”, Neuron 3;84(5):1079–90. doi: 10.1016/j.neuron.2014.10.048. Epub 2014 Nov 11. PMID: 25467986; PMCID: PMC4360978.

[21] A. Aggarwal, 2016, “The sensori-motor model of the hippocampal place cells”, Neurocomputing (2016), 10.1016/j.neucom.2015.12.044i

[22] T. Li, A. Arleo, D. Sheynikhovich, 2020, “Modeling place cells and grid cells in multi-compartment environments: Entorhinal-hippocampal loop as a multisensory integration circuit”, Neural Netw. 2020 Jan;121:37–51. doi: 10.1016/j.neunet.2019.09.002. Epub 2019 Sep 6. PMID: 31526953.

[23] R. M. Grieves, É. Duvelle, E. R. Wood, P. A. Dudchenko, 2017, “Field repetition and local mapping in the hippocampus and the medial entorhinal cortex”, J Neurophysiol. 2017 Aug 16;118(4):2378–2388. doi: 10.1152/jn.00933.2016.

[24] G. Chen, J.A. King, N. Burgess, J. O’Keefe, 2013, “How vision and movement combine in the hippocampal place code”, Proc Natl Acad Sci USA 110:378–383, 2013. doi:10.1073/pnas.1215834110.

[25] M.P. Witter and H.J. Groenewegen, 1984, “Laminar origin and septotemporal distribution of entorhinal and perirhinal projections to the hippocampus in the cat”, J. Comp. Neurol., 224, 371–384.

[26] M.P. Witter, H.J. Groenewegen, F.H. Lopes da Silva and A. H. M. Lohman, 1989, “Functional organisation of the extrinsic and intrinsic circuitry of the parahippocampal region”, Prog. Neurobiol., 33, 161–253.

[27] N.M. Van Strien, N.L. Cappaert, M.P. Witter, 2009, “The anatomy of memory: an interactive overview of the parahippocampal-hippocampal network”, Nature reviews Neuroscience 10, 272–282.

[28] M. P. Witter, et. al., 2009, “Cortico-Hippocampal Communication by Way of Parallel Parahippocampal-Subicular Pathways”, Hippocampus, 10:398–410.

[29] J. J. Peng, B. Throm, M. Najafian Jazi, et al., 2025, “Grid cells accurately track movement during path integration-based navigation despite switching reference frames”, Nat Neurosci 28, 2092–2105. 10.1038/s41593-025-02054-6.

[30] F. Carpenter, D. Manson, K. Jeffery, N. Burgess, C. Barry, 2015, “Grid cells form a global representation of connected environments”, Curr Biol, 25: 1176–1182, 2015. doi:10.1016/j.cub.2015.02.037.

[31] E. Marozzi, L. L. Ginzberg, A. Alenda, K. J. Jeffery, 2015, “Purely translational realignment in grid cell firing patterns following nonmetric context change”, Cereb Cortex 25: 4619–4627, 2015. doi:10.1093/cercor/bhv120.

[32] J. A. Pérez-Escobar, O. Kornienko, P. Latuske, L. Kohler, K. Allen, 2016, “Visual landmarks sharpen grid cell metric and confer context specificity to neurons of the medial entorhinal cortex”, eLife 5: e16937, 2016. doi:10.7554/eLife.16937.

[33] C. Wang, H. Lee, G. Rao, et al., 2024, “Multiplexing of temporal and spatial information in the lateral entorhinal cortex”, Nat Commun 15, 10533 (2024). 10.1038/s41467-024-54932-5

[34] R.J. Gardner, E. Hermansen, M. Pachitariu, et al., 2022, “Toroidal topology of population activity in grid cells”, Nature 602, 123–128. 10.1038/s41586-021-04268-7.

[35] E. I. Moser, E. Kropff, M. B. Moser, 2008, “Place cells, grid cells, and the brain’s spatial representation system”, Annu Rev Neurosci. 31:69–89. doi: 10.1146/annurev.neuro.31.061307.090723. PMID: 18284371.

[36] J. Krupic, M. Bauza, S. Burton, et al., 2015, “Grid cell symmetry is shaped by environmental geometry”, Nature 518, 232–235 (2015). 10.1038/nature14153.

[37] R. J. Low, Y. Gu, D. W. Tank, 2014, “Cellular resolution optical access to brain regions in fissures: imaging medial prefrontal cortex and grid cells in entorhinal cortex”, Proc Natl Acad Sci U S A. 2014 Dec 30;111(52):18739–44. doi: 10.1073/pnas.1421753111. Epub 2014 Dec 12. PMID: 25503366; PMCID: PMC4284609.

[38] J. D. Monaco, L. F. Abbott, 2011, “Modular Realignment of Entorhinal Grid Cell Activity as a Basis for Hippocampal Remapping”, Journal of Neuroscience, 22 June 2011, 31 (25) 9414–9425; 10.1523/JNEUROSCI.1433-11.2011

[39] Alme et. al., 2014, “Place cells in the hippocampus: Eleven maps for eleven rooms”, PNAS:111(52), 18428–18435.

[40] Lian, Burkitt, 2021, “Learning an Efficient Hippocampal Place Map from Entorhinal Inputs Using Non-Negative Sparse Coding”, eNeuro, 8 (4) ENEURO.0557-20.2021.

[41] C. Lykken, B. R. Kanter, A. Nagelhus, M. Guardamagna, J. Carpenter, M.-B. Moser, E. I. Moser, 2024, “Independence of grid cell modules during hippocampal remapping”, Program No. PSTR190.04. Neuroscience Meeting Planner. Chicago, IL. Society for Neuroscience, 2024. Online.

[42] Y. Gronich, V. A. Normand, M.-B. Moser, E. I. Moser, Y. Burak, 2024 “Multi-module grid cells in the medial entorhinal cortex”, Program No. PSTR190.02. Neuroscience Meeting Planner. Chicago, IL. Society for Neuroscience, 2024. Online.

[43] T. Solstad, E. I. Moser, G. T. Einevoll, 2006, “From grid cells to place cells: a mathematical model”, Hippocampus;16(12):1026–31. doi: 10.1002/hipo.20244. PMID: 17094145.

[44] J. O’Keefe, M. L. Recce, 1993, “Phase relationship between hippocampal place units and the EEG theta rhythm”, Hippocampus, 3(3):317–30. doi: 10.1002/hipo.450030307. PMID: 8353611.

